# LiFE, a multimodal circadian intervention, improves sleep, glycemic control, and recognition memory

**DOI:** 10.64898/2026.03.12.711428

**Authors:** Yu Shi, Stephen D. Rozen, Jordan T. Swint, Williams A. McRoberts, Sophia N. McCurry, Ricardo Salinas, Elizabeth G. Moffett, Clara M. Pollock, Lila R. Goldstein, Soraya S. Katzev, Matthew E. Carter, George S. Bloom, Ali D. Güler

**Affiliations:** Departments of Biology, University of Virginia; Charlottesville, Virginia 22904, USA; Department of Biology, Program in Neuroscience, Williams College, Williamstown, MA 01267, USA; Departments of Cell Biology, University of Virginia; Charlottesville, Virginia 22904, USA; Departments of Neuroscience; University of Virginia; Charlottesville, Virginia 22904, USA

**Keywords:** Circadian entrainment, photoperiod, time-restricted feeding, time-restricted exercise, Alzheimer’s disease, sleep, cognition, glucose metabolism, 5xFAD, PS19

## Abstract

In mammals, sleep is regulated by the central circadian system, which responds to environmental timing cues including light, exercise and availability of food. In this study, we developed a light-, food-, and exercise-based daily lifestyle intervention (LiFE) that combines the effects of multiple circadian entrainment cues on central clock function, ultimately strengthening central clock rhythms. In wild-type (WT) mice, LiFE consolidated nocturnal activity, enhanced suprachiasmatic nucleus rhythmicity, and increased sleep time. Despite comparable caloric intake to control conditions, LiFE lowered baseline blood glucose, reduced glycemic variability, and improved glucose tolerance. We found long-term LiFE treatment improved recognition memory in WT mice. Sleep and circadian disruption are commonly observed in patients with Alzheimer’s disease (AD), the most prevalent neurodegenerative disorder. We applied long-term LiFE treatment in two AD mouse models (5xFAD and 5xFAD/PS19). Alongside a subtle reduction in AD histopathology, LiFE produced near-significant trends toward improved motor performance and recognition memory. Together, these findings support multimodal circadian chronotherapy as a non-pharmacological approach in which integrated light, feeding, and exercise entrainment promotes sleep and metabolic health.

## Introduction

The circadian system is evolutionarily conserved and organizes physiology and behavior to adapt to daily environmental cycles. In mammals, cell-autonomous molecular clocks generate transcriptional–translational feedback loops of clock genes that drive ∼24-hour rhythms (1). To generate coherent organism-wide rhythms, the hypothalamic suprachiasmatic nucleus (SCN) functions as the central pacemaker, receiving direct input from intrinsically photosensitive retinal ganglion cells (ipRGCs) to entrain these rhythms to the day/night cycle (2, 3). Although light is the dominant timing cue (zeitgeber) that entrains the circadian clock, other cues, including food intake, physical activity, stress, and ambient temperature cycles, can also entrain circadian rhythms (4–7). These timing cues act on peripheral oscillators including liver and skeleton muscles and influence SCN rhythmicity (8–11).

Because circadian rhythms exert broad control over physiology, aligning daily behaviors with the circadian clock has emerged as a strategy to improve health. Circadian lifestyle interventions primarily leverage the timing of behaviors such as eating and physical activity, along with light-based cues. For instance, time restricted feeding (TRF) during the active phase enhances rhythmic gene expression and improves metabolic function in obese mice (12, 13). In humans, intermittent fasting improves glucose tolerance and cardiovascular health (14). Scheduled exercise late in the active phase has also been shown to improve both molecular and behavioral rhythms in a clock deficient mouse model (15). Light based interventions such as short photoperiods promote synchrony of SCN neurons and increase central clock rhythm amplitude (16). Although exposure to light at night is widely recognized as detrimental to circadian and metabolic health, the therapeutic potential of short photoperiods to improve overall health, particularly in disease models, remains largely unexplored. Moreover, despite the expectation that multiple entrainment cues act synergistically, most studies test a single zeitgeber in isolation. As a result, it remains poorly understood whether combining photic and nonphotic entrainment cues produces additive or synergistic physiological benefits.

Alzheimer’s disease (AD) is the most prevalent neurodegenerative disease with no effective treatment, and an estimated total cost of $300-400 billion in the United States annually (17). Studies in recent years have identified a bidirectional relationship between circadian disruption and AD pathology, suggesting that circadian dysfunction may actively contribute to disease progression (18). Typical sleep disruptions, including increased daytime sleep and sleep fragmentation, are commonly observed in AD patients(19). Recent studies identified alterations in circadian gene expression in human AD samples (20). Furthermore, circadian misalignment disrupts sleep and metabolic homeostasis, and contributes to cognitive decline and neurodegeneration (21). Circadian treatments that target feeding schedules, such as intermittent fasting and TRF, are emerging as promising strategies for the care of AD patients, and have been shown to ameliorate AD pathology and improve cognition in mouse models (22) (23).

To determine how combined entrainment cues enhances physiological benefits, we developed a multimodal circadian entrainment paradigm using time-restricted <u>li</u>ght, food, and <u>e</u>xercise (LiFE). We showed that LiFE treatment consolidates diurnal activity and enhances SCN rhythmicity, increases sleep time and decreases blood glucose levels in wild-type (WT) mice. In addition, we found short photoperiod alone did not improve glucose metabolism and increased REM sleep fragmentation. Using two aggressive AD mouse models, we found that long-term LiFE entrainment produced modest, directionally consistent benefits across functional and pathological readouts. In 5xFAD mice, LiFE was associated with improved task acquisition and partial amelioration of motor deficits. In the hybrid 5xFAD/PS19 model, LiFE showed trends toward improved novel object recognition and reduced hippocampal AD pathology. Although these effects did not reach statistical significance, their convergence across two stringent models supported the potential of multimodal chronotherapeutic strategies. Overall, our data suggested that coordinated integration of multiple circadian entrainment cues, rather than single zeitgebers alone, promoted physiological health in both WT and AD mice and warranted further evaluation in broader and more clinically representative contexts.

## Results

### LiFE entrainment consolidates diurnal activity and enhances SCN rhythmicity in WT mice

To investigate the roles of light, feeding, and exercise entrainment individually and in combination, we designed a panel of circadian entrainment regimens. We assessed circadian behavioral outputs by monitoring beam-break locomotor activity in wild-type (WT) mice under each condition (Fig. 1a-j). Control and time-restricted light (TRL) groups had ad libitum food and wheel access on 12:12-h or 8:16-h light–dark (LD) cycles, respectively. In both groups, activity onset coincided with lights-off and total activity did not differ (Fig. 1a, f). Using TRL (8:16 light–dark cycle) as the reference condition, we next introduced either time-restricted feeding (TRFL) during the first 8 hours of the dark phase or time-restricted exercise (TREL), with running wheels available starting 8 hours after lights-off. TRFL did not change the overall activity pattern but modestly increased activity in the 2 hours preceding food availability, consistent with food-anticipatory activity (FAA) (control: 268 ± 106; TRFL: 1278 ± 459 beam breaks; mean ± SEM)(Fig. 1c, g, j). In the time-restricted exercise schedule, although wheels were available during the light phase for logistical reasons, no daytime running was detected (Figure. S1c). TREL mice initiated general locomotor activity upon wheel availability (8 h after dark onset) rather than at lights-off, showing suppressed early-night activity and a compressed active phase (∼8 h vs ∼12 h in control and TRL; Fig. 1d, g). Finally, we implemented a schedule with simultaneous restriction of <u>li</u>ght, food, and <u>e</u>xercise (LiFE). LiFE mice showed food-anticipatory activity and a shifted activity onset, consistent with combined food and exercise entrainment as observed in TRFL and TREL conditions independently (Fig. 1e, h). Notably, relative to controls, the locomotor activity changes induced by TRFL or TREL were further accentuated under the LiFE schedule, leading to a significant reduction in total daily ambulation in LiFE mice (Fig. 1i). In addition, FAA was significantly increased in LiFE-treated mice relative to TRL and control groups (Fig. 1j). Together, LiFE treated mice exhibited strong food entrainment behavior and had consolidated active phase compared to control mice.

**Figure 1:**
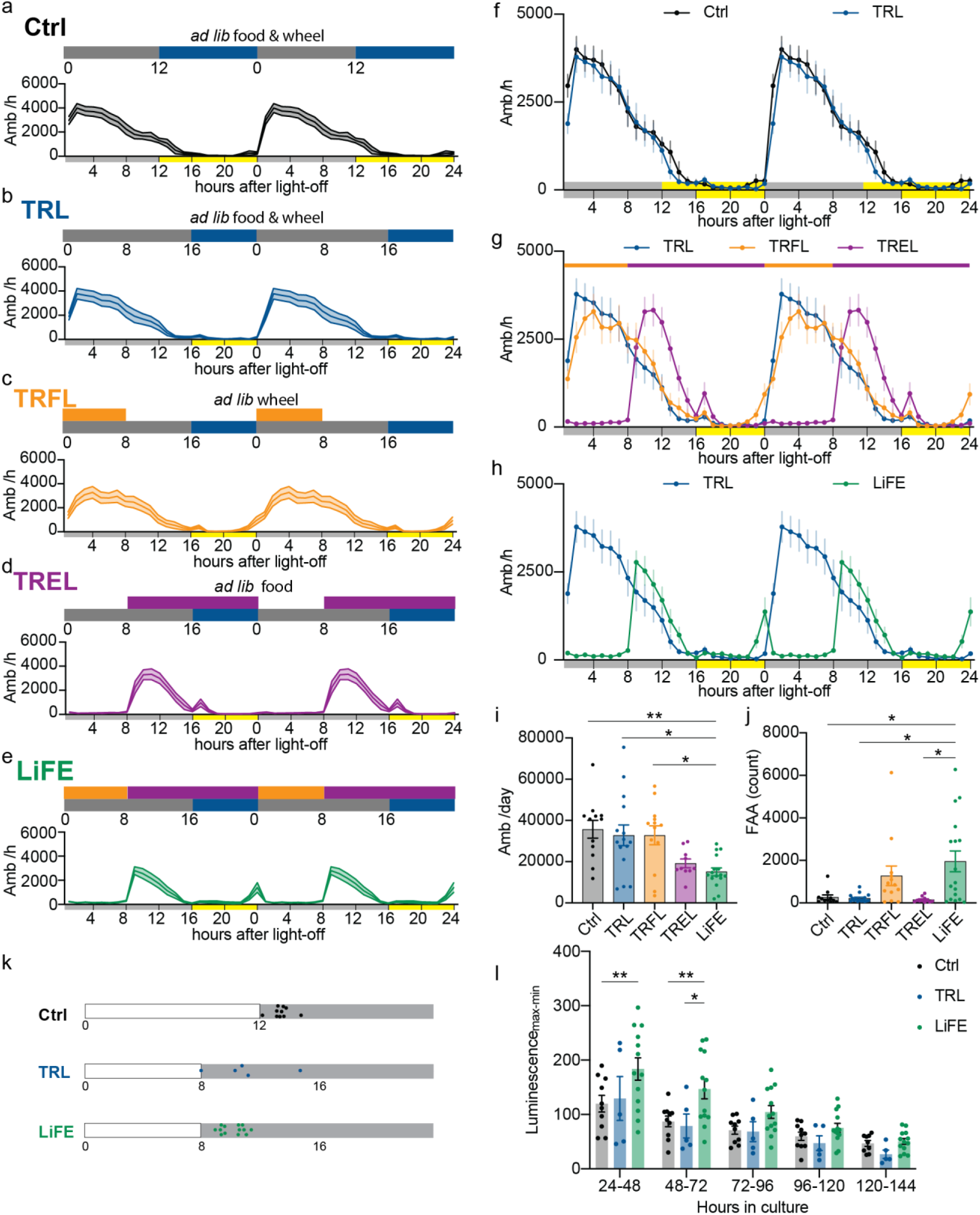
Time-restricted light, food and exercise altered diurnal activity and enhanced central clock amplitude. **a-e)** Schematics of treatment schedules and double-plotted ambulatory activity (Amb, count/hour) of WT mice. Control (n = 12), time-restricted light (TRL, n = 15), time-restricted feeding and light (TRFL, n = 13), time-restricted exercise and light (TREL, n = 10), time-restricted light, feeding and exercise (LiFE, n = 16). Blue bar: light exposure; grey bar: dark; orange bar: access to food; purple bar: access to running wheels. Numbers indicate Zeitgeber time (ZT, hours after light on). **f)** Comparison of ambulatory activity between control and TRL, yellow and grey bars on the x axis indicate light and dark. **g)** Comparison of ambulatory activity of TRL, TRFL and TREL. Orange and purple bars indicate access to food and wheels, yellow and grey bars on the x axis indicate light and dark. **h)** Comparison of ambulatory activity of TRL and LiFE. **i)** Daily total ambulatory activity counts (one-way ANOVA with Bonferroni correction, \**P*<0.05, \*\**P*<0.01) **j)** Food anticipatory activity (total activity during 2 hours before light-off) (Browns-Forsythe test with Dunnett’s T3 correction, \**P*<0.05). **k)** illustrations showing ZT times(colored dots) of peak values measured in *Per2* luminescence, blank bars indicate light phase, grey bars indicate dark phase. **l)** Amplitude of *Per2* luminescence in SCN cultures from control(n = 10), TRL (n = 5) and LiFE (n = 13) treated mice(mixed-effect model with Bonferroni correction, \**P*<0.05; \*\**P*<0.01). Data are represented as means ± SEM; see statistical details in the method section.

To determine whether LiFE enhances the rhythmicity of the central circadian oscillator, we prepared organotypic SCN slices from Per2::Luc reporter mice, in which luciferase expression is driven by the Per2 promoter, and monitored bioluminescence as a real-time readout of Per2 rhythms. Per2::Luc mice were habituated to control, TRL, or LiFE schedules for 2 weeks prior to tissue collection. In culture, the peak phase of bioluminescence rhythms aligned with dark onset across all groups (Fig. 1k). However, SCN slices from LiFE mice displayed an approximately 50% higher rhythm amplitude at 24 to 48 h compared with control and TRL groups (Fig. 1l). This elevated amplitude persisted beyond 72 h in vitro, indicating that LiFE strengthens SCN rhythmicity in a manner that outlasts the entrainment cues.

### Food and exercise entrainment increase sleep time and mitigate fragmentation under short photoperiod in WT mice

The circadian system plays a central role in regulating sleep timing and architecture. To determine how LiFE treatment affects sleep duration and quality, we recorded electroencephalogram (EEG) activity in WT mice habituated to control, TRL, or LiFE schedules. As shown previously, control mice slept predominantly during the light phase with a smaller late-night bout (24). TRL modestly advanced both the offset of the major sleep bout and the onset of the late-night bout (Fig. 2a). In contrast, LiFE mice exhibited a marked increase in early-night sleep that coincided with reduced locomotion, resulting in ∼30% more total dark-phase sleep compared to TRL mice (Fig. 2b,d). Light-phase sleep was also ∼9% higher in LiFE mice than controls, and total 24-h total sleep was increased significantly relative to TRL mice but not control mice (Fig. 2c, e). NREM sleep, which comprises over 70% of total mouse sleep, mirrored these sleep patterns, although the increase in light-phase NREM sleep did not reach statistical significance (Fig. 2f-j). The diurnal distribution of REM sleep was not changed by either TRL or LiFE schedules, however, the total 24-h REM sleep was reduced in TRL mice (Fig. 2k-m). This reduction was not specific to either the dark or light phase and was partially rescued by the addition of food and exercise entrainment in the LiFE schedule (Fig. 2m-o). We did not observe differences in microawakenings, stage transitions, or NREM sleep bout length across conditions (Fig. 2p-r). However, REM bout length was shortened under short photoperiod conditions and was not fully restored by LiFE, particularly during the light phase (Fig. 2s-u). In conclusion, these results indicate that food and exercise entrainment in the LiFE schedule increases NREM sleep during both the dark and light phases and partially compensates for sleep fragmentation induced by a short photoperiod.

**Figure 2:**
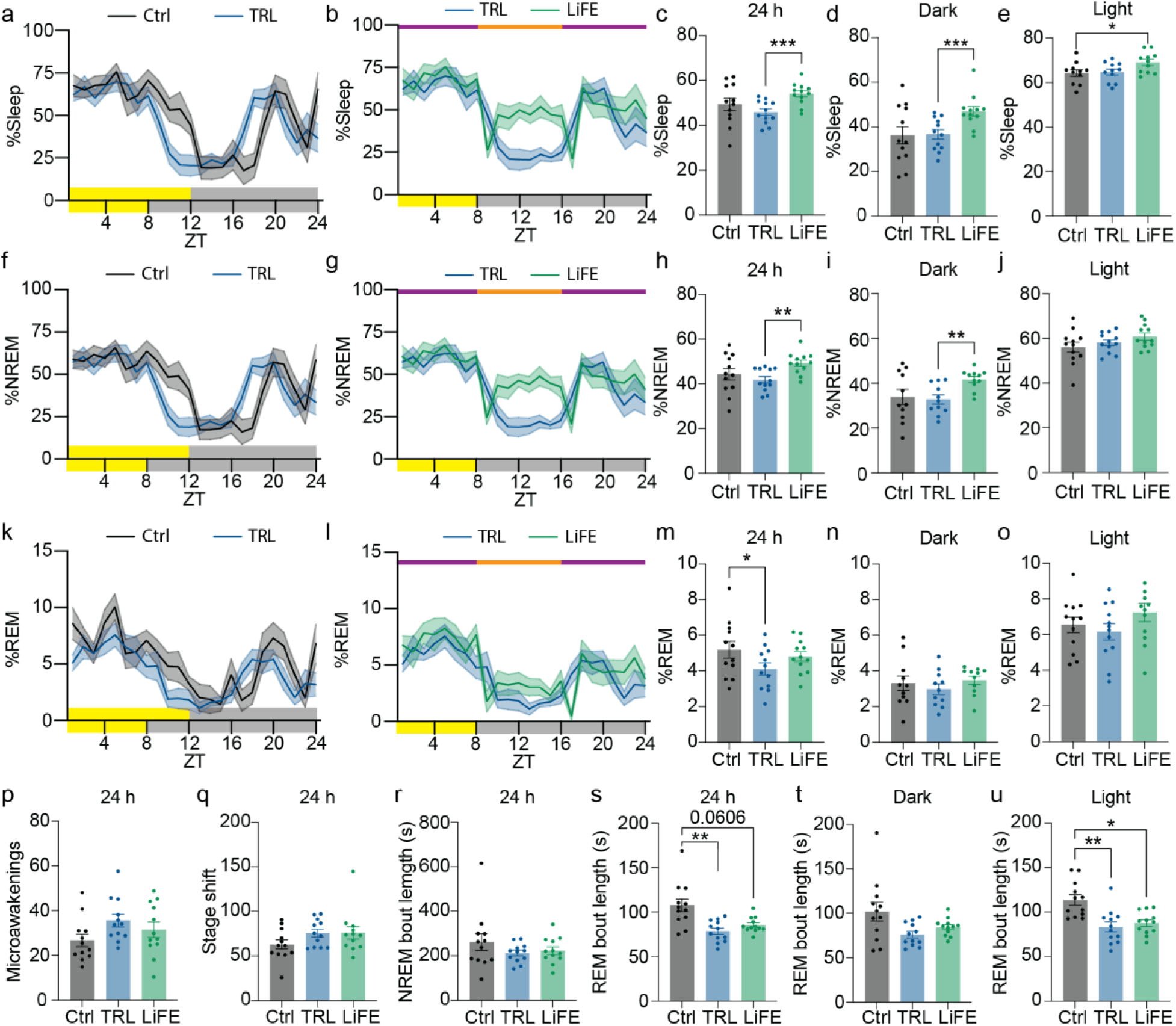
Effects of LiFE treatment on sleep time and structure. **a)** Percentage of total sleep in electroencephalogram (EEG) recordings in WT mice (n =12) habituated to control and TRL schedule in 1h bin. Yellow and grey bars on the x axis indicate light and dark. **b)** Percentage of total sleep in EEG recordings in WT mice (n =12) habituated to TRL and LiFE schedule in 1h bin. Orange and purple bars indicate access to food and wheels. **c)** Calculations of average daily sleep percentage (paired one-way ANOVA with Bonferroni correction, \*\*\**P*<0.001). **d)** Calculations of average sleep percentage in the dark phase (paired one-way ANOVA with Bonferroni correction, \*\*\**P*<0.001). **e)** Calculations of average sleep percentage in the light phase (paired one-way ANOVA with Bonferroni correction, \**P*<0.05). **f)** Percentage of NREM sleep in EEG recordings in WT mice (n =12) habituated to control and TRL schedule in 1h bin. **g)** Percentage of NREM sleep in EEG recordings in WT mice (n =12) habituated to TRL and LiFE schedule in 1h bin. **h)** Calculations of average daily NREM sleep percentage (paired one-way ANOVA with Bonferroni correction, \*\**P*<0.01). **i)** Calculations of average NREM sleep percentage in the dark phase (paired one-way ANOVA with Bonferroni correction, \*\**P*<0.01). **j)** Calculations of average NREM sleep percentage in the light phase. **k)** Percentage of REM sleep in EEG recordings in WT mice (n =12) habituated to control and TRL schedule in 1h bin. **l)** Percentage of REM sleep in EEG recordings in WT mice (n =12) habituated to TRL and LiFE schedule in 1h bin. **m)** Calculations of average daily REM sleep percentage (paired one-way ANOVA with Bonferroni correction, \**P*<0.05). **n)** Calculations of average REM sleep percentage in the dark phase. **o)** Calculations of average REM sleep percentage in the light phase. **p)** Counts of microawakenings in control, TRL and LiFE treated WT mice. **q)** Counts of stage shift in control, TRL and LiFE treated WT mice. **r)** Average bout lengths of NREM sleep in control, TRL and LiFE treated WT mice. **s)** Average bout lengths of REM sleep in control, TRL and LiFE treated WT mice (paired one-way ANOVA with Bonferroni correction, \*\**P*<0.01). **t)** Average bout lengths of REM sleep in the dark phase. **u)** Average bout lengths of REM sleep in the light phase (paired one-way ANOVA with Bonferroni correction, \**P*<0.05, \*\**P*<0.01). Data are represented as means ± SEM; see statistical details in the method section.

### LiFE reduces baseline blood glucose and stabilizes glucose dynamics

Improved metabolic health, particularly glycemic control, is a well-established effect of time-restricted feeding (25). Therefore, we examined how LiFE treatment affected feeding and glucose dynamics. Daily food intake and body weight were comparable across control, TRL, and LiFE groups, indicating that any differences in metabolic readouts would be independent of caloric intake or weight change. (Fig. 3a,b). To quantify daily feeding patterns, we measured food intake at 4-hour intervals. As expected from the imposed time restriction, LiFE mice consumed all their food within an 8-h window, whereas TRL mice distributed feeding across ∼16 h of the dark phase (Fig. 3c). Next, we investigated how LiFE treatment affects baseline blood glucose over 24 h by measuring levels every 4 hours. Relative to control (157.8 ± 3.86 mg/dL) and TRL (147.5 ± 3.55 mg/dL), LiFE mice exhibited lower average blood glucose (137.2 ± 2.72 mg/dL) and reduced glucose variability, measured as the difference between the maximum and minimum glucose levels (Fig. 3d-f). In glucose tolerance tests conducted 4 h after dark onset, when baseline differences were greatest, LiFE mice showed improved glucose clearance relative to TRL (Fig. 3g,h). Together, these data indicate that LiFE enhances glucose metabolism by lowering baseline glucose and reducing daily fluctuations.

**Figure 3:**
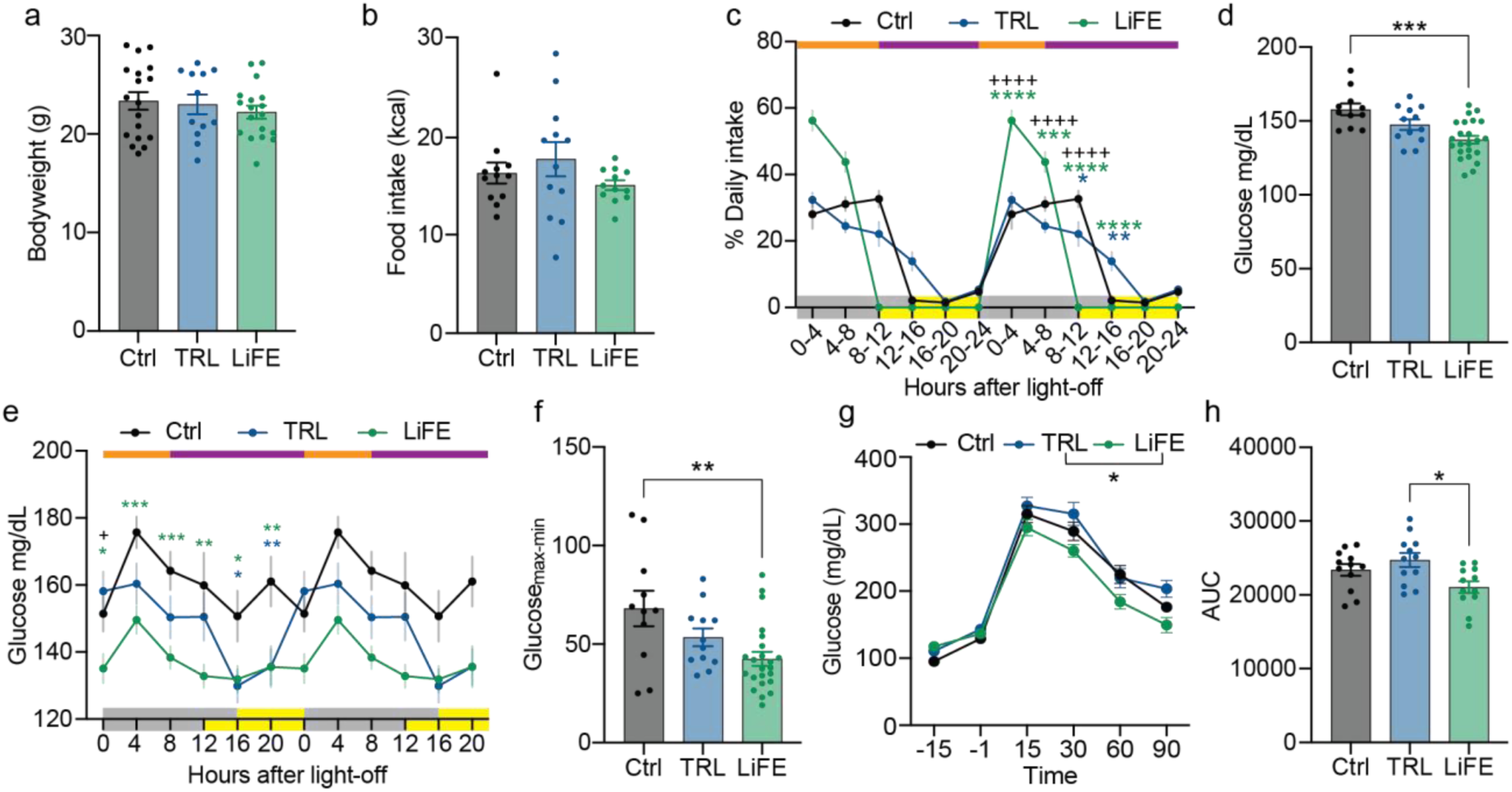
LiFE treatment reduced baseline blood glucose level and fluctuation. **a)** Bodyweight of WT mice on control (n = 18), TRL (n = 12) and LiFE (n = 18). **b)** Daily food consumption of WT mice (n = 12) on control, TRL and LiFE schedule. **c)** Food consumption percentage of WT mice on control (n = 6), TRL (n = 12) and LiFE (n = 18) schedules in a 4 hour bin. Orange and purple bars indicate access to food and wheels, yellow and grey bars on the x axis indicate light and dark. (mixed-effect model with Bonferroni correction, *P<0.05, **P<0.01, ***P<0.001, ****P<0.001, green stars: LiFE vs. control; blue stars: TRL vs. control; +: LiFE vs. TRL). **d)** Averaged daily baseline blood glucose calculated from e) (one-way ANOVA with Bonferroni correction, ***P<0.001). **e)** Baseline blood glucose of WT mice on control (n = 11 at time 12, 16, 20; n = 23 at time 0, 4, 8), TRL (n = 12) and LiFE (n = 23) schedule (mixed-effect model with Bonferroni correction, *P<0.05, **P<0.01, ***P<0.001). **f)** Difference between maximum and minimum baseline blood glucose calculated from e) (one-way ANOVA with Bonferroni correction, **P<0.01). **g)** Glucose tolerance test of WT mice (n = 12) on control, TRL and LiFE schedule at dark + 4 hours (mixed-effect model with Bonferroni correction, *P<0.05). **h)** Area under curve calculated from g) (one-way ANOVA with Bonferroni correction, *P<0.05). Data are represented as means ± SEM; see statistical details in the method section.

### LiFE treatment improved recognition memory in WT mice and mitigated disease-associated behaviors in AD mouse models

Sleep plays a vital role in memory consolidation, and circadian disruption can impair learning and memory (26, 27).To assess whether LiFE influences cognition, we applied long-term LiFE treatment to WT mice from 2 to 3 months of age until 7 to 8 months (Fig. 4a). We then evaluated hippocampal-dependent spatial and recognition memory using the Morris water maze (MWM) and novel object recognition (NOR) tests, respectively. In the MWM, mice learn to locate a hidden escape platform using distal visual cues. On day 1, animals underwent habituation with a visible platform to assess baseline performance, including motor ability and motivation, whereas subsequent acquisition and probe trials assessed spatial learning and memory. LiFE treatment did not alter MWM performance in WT mice, likely reflecting a ceiling effect as all groups performed at a high level (Fig. 4b–e). In contrast, in the NOR task with a 24-hour delay between familiarization and testing, LiFE-treated WT mice showed significantly higher recognition scores than controls (Fig. 4f).

**Figure 4:**
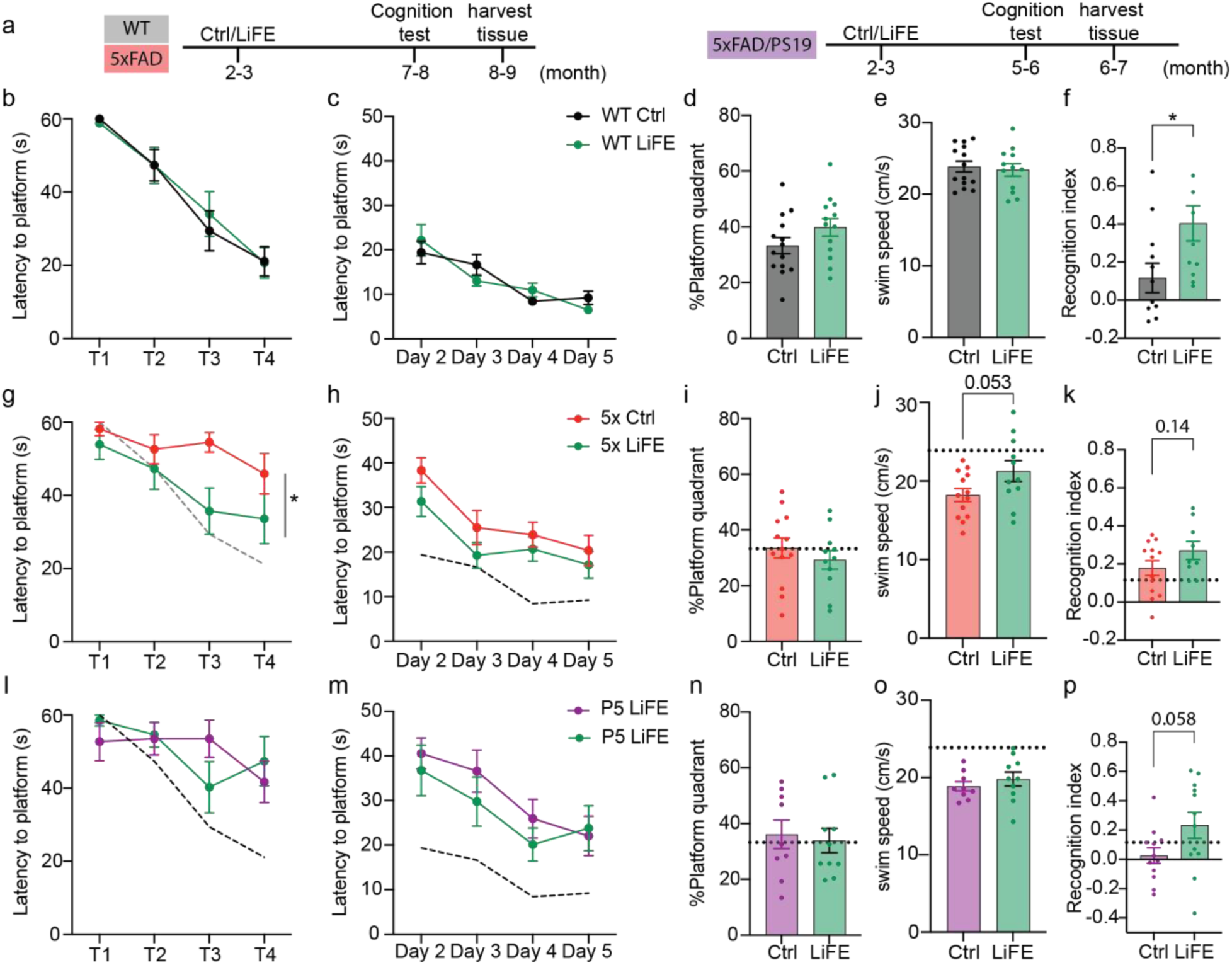
Long-term LiFE treatment improves learning in Morris water maze and novel object recognition. **a)** Schematics of experiment timeline of WT, 5xFAD and 5xFAD/PS19 mice with long-term LiFE treatment. **b)** Latency to platform on day 1 measured in MWM in WT mice (control: n = 14; LiFE: n = 13). **c)** Latency to platform on acquisition days in MWM in WT mice. **d)** Percentage of time spent in target quadrant on probe day of MWM in WT mice. **e)** Swim speed of WT mice in MWM. **f)** Recognition index (see method) of WT mice in NOR (control: n = 12; LiFE: n = 11, Student’s t-test, \**P*<0.05). **g)** Latency to platform on day 1 in MWM in 5xFAD mice (control: n = 13; LiFE: n = 11). The dashed line indicates the level of WT controls. (mixed-effect model with Bonferroni correction, \**P*<0.05) **h)** Latency to platform on acquisition days in MWM in 5xFAD mice. **i)** Percentage of time spent in target quadrant on probe day of MWM in 5xFAD mice. **j)** Swim speed of 5xFAD mice in MWM (Student’s t-test). **k)** Recognition index of 5xFAD mice in NOR(control: n = 13; LiFE: n = 9, Student’s t-test). **l)** Latency to platform on day 1 in MWM in 5xFAD/PS19 mice (n = 10). **m)** Latency to platform on acquisition days in MWM in 5xFAD/PS19 mice. **n)** Percentage of time spent in target quadrant on probe day of MWM in 5xFAD/PS19 mice. **o)** Swim speed of 5xFAD/PS19 mice in MWM. **p)** Recognition index of 5xFAD/PS19 mice in NOR(n = 12, Student’s t-test). Data are represented as means ± SEM; see statistical details in the method section.

Circadian and sleep disruptions are both risk factors and preclinical symptoms of Alzheimer’s disease, and in healthy individuals, sleep deprivation leads to decline of learning and memory (28, 29). In addition, diabetes is a common comorbidity in AD. Because our WT data show that LiFE increases sleep time while improving glycemic control and recognition memory, we hypothesized that long-term LiFE treatment might ameliorate AD-related phenotypes. To test this, we implemented long-term LiFE in two AD mouse models, aligning the intervention window to each model’s disease progression (Fig. 4a). We began with 5xFAD, a widely used model of rapid amyloid plaque accumulation (Fig. 4g–k). 5xFAD mice overexpress human APP carrying the Swedish (K670N/M671L), Florida (I716V), and London (V717I) mutations together with PSEN1 mutations (M146L and L286V). Because 5xFAD robustly models amyloid pathology but lacks tauopathy, we complemented it with a second model that incorporates both hallmark lesions. PS19 mice overexpress human tau carrying the P301S mutation, and prior work crossed PS19 with 5xFAD to generate the hybrid 5xFAD/PS19 line (hereafter 5P), which develops both amyloid and tau pathology (30). Relative to traditional mixed AD models, 5P progresses rapidly while retaining a broad spectrum of AD-relevant pathological features. We therefore selected 5P as the second model in our study (Fig. 4l–p). LiFE treatment was initiated at the onset of amyloid plaque deposition at 2.5 months (31). Cognitive tasks were performed at 7.5 months of age, after the age at which impairment has been reported in 5xFAD mice (32), and at 5.5 months of age in 5P mice, reflecting this model’s more rapid and severe disease progression. We first assessed hippocampal-dependent spatial learning and memory using MWM, which also provides a sensitive readout of motor performance in these models. During the visible-platform day (day 1), 5xFAD control mice showed longer escape latencies than WT mice (Fig. S2a). LiFE-treated 5xFAD mice reached the platform faster than controls on this day (Fig. 4g), consistent with improved task performance under conditions that minimize spatial memory demands. However, LiFE did not produce significant differences during hidden-platform acquisition or the probe trial (Fig. 4h, i). Because motor impairment is a reported phenotype in 5xFAD mice (33), we also quantified swimming speed and detected a motor deficit in our cohort (Fig. S2b). LiFE treatment was associated with a trend toward faster swim speed in 5xFAD mice (control: 18.21 ± 0.82 cm/s; LiFE: 21.27 ± 1.31 cm/s; Fig. 4j). Similarly, 5P mice exhibited learning and motor deficits relative to WT in MWM (Fig. S2a, b), but LiFE-treated and control 5P mice performed comparably across all tested days (Fig. 4l–o). We next evaluated recognition memory using the novel object recognition (NOR) task. Relative to WT, 5xFAD control mice did not show a detectable recognition memory deficit, whereas 5P control mice exhibited clear impairment (Fig. S2e). Consistent with this baseline profile, LiFE did not significantly increase the recognition index in 5xFAD mice, although values trended upward (control: 0.18 ± 0.04; LiFE: 0.27 ± 0.05; Fig. 4k). In 5P mice, LiFE treatment produced a trend toward improved recognition memory (recognition index: 0.23 ± 0.09 vs. 0.03 ± 0.05 in controls), but high variability prevented statistical significance (Fig. 4p). Together, these results indicate that long-term LiFE robustly improved recognition memory in WT mice and produced subtle, model-dependent benefits across two AD mouse models.

### LiFE treatment modestly altered AD pathology in the hippocampus of 5P mice

Finally, we quantified neuropathology in 5xFAD and 5P mice with and without long-term LiFE treatment. Using immunohistochemistry, we assessed amyloid-β related deposition with the D54D2 antibody, which recognizes multiple Aβ species as well as full-length APP and APP fragments containing the Aβ sequence, and robustly labels plaques in vivo. We assessed tau pathology using AT8, which detects abnormally phosphorylated tau at pSer202/pThr205. At the time point of tissue collection, the hippocampus and cerebral cortex are the primary regions exhibiting amyloid and tau pathology in 5xFAD and 5P mice (30, 34, 35). In the hippocampus, we quantified pathology staining in the CA1 and dentate gyrus separately. In the cerebral cortex, we analyzed staining in the medial region, specifically retrosplenial cortex (RS), and in the lateral region including perirhinal and entorhinal cortex. In 5xFAD mice, control and LiFE-treated mice had equal fractions of areas of D54D2 staining in all brain regions we measured (Fig. 5 a,b). In LiFE-treated 5xFAD/PS19 mice, we observed a trend toward reduced D54D2 and AT8 staining in the hippocampus (Fig. 5c–e). In the dentate gyrus, mean AT8 staining was similar between groups, but the distribution shifted, with more LiFE-treated mice showing lower staining (Fig. 5e). In contrast, amyloid and tau pathology in cortical regions was unchanged (Fig. 5c–e). Overall, in this hybrid model, long-term LiFE treatment was associated with a modest, hippocampus-specific trend toward reduced AD pathology.

**Figure 5:**
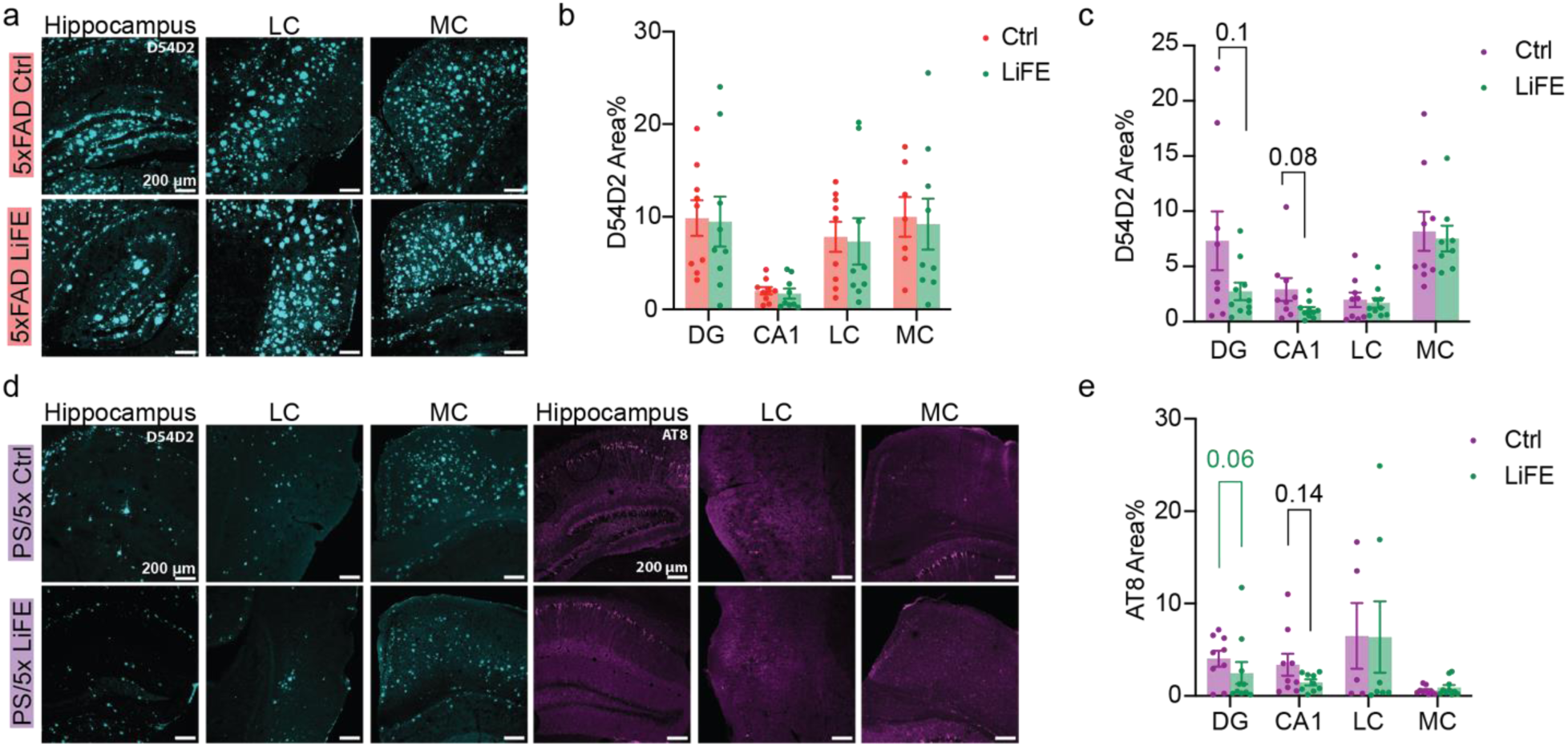
Long-term LiFE treatment reduced AD pathology in the hippocampus. **a)** Representative images of D54D2 immunohistochemistry (IHC) staining in 5xFAD mice, 10X objective lens. **b)** Quantification of area fractions of D54D2 staining in 5xFAD mice (n = 9). **c)** Quantification of area fractions of D54D2 staining in 5xFAD/PS19 mice (control: n = 9, LiFE: n = 10 DG, CA1, LC, n = 8 MC; Student’s t-test). **d)** Representative images of D54D2 and AT8 IHC staining in 5xFAD/PS19 mice, 10X objective lens. **e)** Quantification of area fractions of AT8 staining in 5xFAD/PS19 mice (control: n = 9 DG, CA1, MC, n = 5 LC; LiFE: n = 10 DG, n = 9 CA1, n = 7 LC, n = 10 MC; black p-value: Student’s t-test; green p-value: Kolmogorov–Smirnov test). Data are represented as means ± SEM; see statistical details in the method section.

## Discussion

Chronotherapy approaches that leverage circadian entrainment to promote health and mitigate disease progression have gained increasing attention in recent years (36). Both scheduled feeding and voluntary exercise have individually shown beneficial effects on physiology and behavior in rodents and humans (15, 37, 38). Notably, several studies have coupled exercise with time-restricted feeding, but timed voluntary exercise specifically designed to enhance circadian entrainment has not been implemented as a therapy for neurodegenerative disease (39, 40). In this study, we aimed to determine whether combining multiple entrainment strategies could amplify their therapeutic potential. We developed a composite entrainment regimen, LiFE (Light, Feeding, and Exercise), that integrates a short photoperiod with time-restricted feeding and scheduled voluntary exercise. In alignment with current understanding of molecular clock mechanisms and nonphotic entrainment, previous work suggests that time-restricted exercise most effectively enhances circadian rhythmicity when applied during the late dark phase (41,15, 37), whereas time-restricted feeding is most commonly administered, and often most effective, during the early dark phase (42). We systematically characterized the outcomes of LiFE treatment, ranging from locomotor activity and sleep to metabolic and cognitive performance, and applied this approach to an Alzheimer’s disease mouse model.

Sleep naturally deteriorates with age, a major risk factor for AD. Abnormal sleep, including reductions in both NREM and REM sleep and shifts in EEG power spectra, has been reported in several AD mouse models, including 5xFAD (43) and PS19 (44). Our EEG analyses revealed that LiFE increased total NREM sleep time during both the light and dark phases, without compromising sleep quality as measured by fragmentation indices. Interestingly, the most prominent increase occurred during the active (dark) phase and coincided with reduced locomotor activity. A similar pattern has been reported in studies where time-restricted voluntary exercise under constant darkness reduced wakefulness in the hours preceding wheel access (4). Some studies interpret increased sleep during the active phase as abnormal (45). In our case, however, the additional sleep was temporally confined and did not disrupt daytime sleep architecture. Thus, the early-night sleep episode we observed may represent an adaptive response to time-restricted feeding or exercise rather than sleep fragmentation. Although this increase occurred at an atypical circadian time, it could still support sleep-dependent clearance processes and influence the rhythmic fluctuation of amyloid-β in interstitial fluid (46), warranting further investigation.

Previous work has shown that time-restricted feeding (TRF) benefits glucose metabolism in obese and aged mouse models (47, 48). Consistent with these reports, we find that comparable metabolic benefits are observed in LiFE-treated healthy WT mice. Notably, LiFE was also associated with increased sleep duration, and sleep is a key contributor to glucose homeostasis (49). Thus, the changes in glucose dynamics observed here likely reflect the combined effects of feeding timing and exercise-based entrainment rather than TRF alone. In addition, we evaluated the contribution of a short photoperiod in WT mice. Short photoperiods can enhance neuronal synchrony within the SCN at cellular and molecular levels (50, 51), but their potential as a therapeutic circadian intervention remains relatively understudied. In our hands, short photoperiod alone did not increase total sleep or improve glycemic regulation and was associated with greater REM sleep fragmentation. Together, these results suggest that light manipulation by itself is insufficient to reproduce the integrated benefits of LiFE.

Overall, LiFE produced selective and consistent improvements in sleep, metabolism, and recognition memory in WT mice, but only subtle trends toward rescuing cognitive deficits or pathology in two AD models. Several factors may have limited the apparent treatment effects in the AD cohorts. First, control animals had access to unlimited voluntary exercise, which has been reported to ameliorate AD-associated phenotypes (52, 53). This may have improved baseline health in the control groups, consistent with our behavioral results: neither 5xFAD nor 5P mice showed clear deficits on the MWM probe trial, and 5xFAD mice exhibited intact recognition memory. Second, 5xFAD may be a less favorable model for circadian chronotherapy because circadian dysfunction is not a prominent or consistent feature. Moreover, circadian phenotypes in 5P mice have not been well characterized in the literature. Although circadian interventions can reverse abnormal diurnal behaviors in other AD models (23, 52, 53), both 5xFAD and 5P displayed only modest differences in locomotor activity relative to WT during the LiFE treatment window (54). Although changes in clock gene expression have been reported in young 5xFAD mice (55), other evidence suggests that the core clock remains rhythmic (56). In our study, LiFE increased SCN Per2 rhythm amplitude, but the extent to which strengthening central clock rhythmicity influences AD pathology remains an open question (56, 57). Finally, substantial inter-individual variability in these models likely reduced our power to detect modest treatment effects. Future studies with larger sample sizes will be important to clarify the magnitude and reproducibility of the directionally consistent trends observed here.

Nevertheless, our study provides a comprehensive framework for integrating light, feeding, and exercise entrainment as a unified circadian intervention. In wild-type mice, LiFE produced clear and consistent benefits across multiple domains, including strengthened SCN rhythmicity, increased sleep, improved glycemic control, and enhanced recognition memory. In two aggressive AD models, LiFE effects were more selective, with directionally consistent trends in specific behaviors and brain regions. Across models, recognition memory showed varying degrees of improvement, and in 5xFAD/PS19 mice the most notable pathological signal was a modest, hippocampus-focused shift with comparatively little change in cortical regions. Together, these findings support multimodal circadian chronotherapy as a tractable, nonpharmacological strategy and motivate continued efforts to define how distinct entrainment cues interact and engage relevant neural and peripheral pathways. An important remaining question is how the timing of intervention onset influences outcomes and whether comparable benefits can be achieved across different stages of disease progression.

## Material and Methods

### Mouse lines

All experiments were carried out in compliance with the Association for Assessment of Laboratory Animal Care policies and approved by the University of Virginia Animal Care and Use Committee. Mice were housed on a 12:12-hour light/dark (LD) cycle with food (PicoLab Rodent Diet 5053) and water *ad libitum* unless otherwise indicated. For experiments, we used 8-week or older male and female C57BL/6J mice, mPER2^luc^ (JAX stock #006852), 5xFAD mice (JAX stock# 034848). 5xFAD/PS19 mice were generated by crossing 5xFAD and PS19 (JAX stock #024841) heterozygous mice.

### Circadian entrainment schedules

#### Short-term entrainment

Short -term entrainment schedule was applied to general activity recording, sleep recording, food intake and bodyweight measurement, glucose tolerance test and baseline glucose measurement in WT mice. Animals were single-housed in running-wheel cages and habituated to LD12:12 (control) or LD8:16 (TRL, TREL, TRFL, LiFE) light-dark cycle in light-tight boxes (∼400 lux fluorescent light illumination). After light cycle habituation, mice with time-restricted feeding treatment (TRFL, LiFE) had *ad libitum* standard food (4 whole pellets) provided to the cage top at light-off time. 8 hours after light-off, remaining food was removed from the top. At the light-off time, mice with time-restricted running wheels (TREL, LiFE) had wheels locked by ties manually, and wheels were unlocked 8 hours after light-off. Body weights were tracked daily in the first week of time-restricted feeding to ensure health of the animals. Experiments were conducted >10 days after the entrainment schedules started.

#### Long-term entrainment

Long-term entrainment time line was applied to WT, 5xFAD and 5xFAD/PS19 mice in cognition tests and histology analysis. Animals were housed on entrainment schedules described above starting from 2-3 months age for >3 months until the end of all experiments.

### General activity recording

Mice were individually housed in transparent wheel-running cages (Nalgene), with IR beam interruption chambers (Columbus Instruments) in light-tight boxes. IR beam interruptions were recorded by Columbus software with a 10 second bin and exported as csv files. Data was organized and binned by 1 hour in RStudio.

### Wheel running recording

Mice were housed in running wheel cages with magnets attached to the wheel. Hall effect sensors were installed on the cage wall to count the cycles of wheel running. Data was recorded and exported by Actimetrics software.

### Lumicycle

mPER2^luc^ mice were individually housed in light-tight boxes on control, TRL or LiFE schedules described above for >2 weeks. Mice were sacrificed between 0-8 hours after light-off in dark. Brains were immediately harvested and dropped into ice cold Hanks’ Balanced Salt Solution (HBSS). After a quick rinse in HBSS, brains were embedded in low melting point agarose (Precisionary Instruments, Natick, MA) and sectioned at 300 micron on a compresstome VF-200 vibrating microtome (Precisionary Instruments, Natick, MA). A floating cell culture insert (0.4μm, 30mm diameter, Millicell) was placed in a 35 mm culture dish containing 1 ml of DMEM medium (D5030, Sigma) supplemented with 3.5 g/L D-glucose, 2 mM Glutamax (Gibico), 10 mM HEPES, 25 U/ml penicillin/streptomycin, 2% B-27 Plus (Gibco), and 0.1 mM D-Luciferin sodium salt (Tocris). Brain regions containing SCN were dissected and placed on the center of cell culture insert one section per dish. The culture dishes were covered with 40 mm diameter glass cover slides and sealed with high-vacuum grease (Dow Corning), and maintained in a non-humidified incubator at 36.8 ℃. Bioluminescence was recorded in 10 min intervals by a 32-channel/4-photomultiplier tube luminometer LumiCycle (Actimetrics) in the incubator. Bioluminescence data were detrended in LumiCycle Analysis software (Actimetrics) and exported as csv files. Data from the first 24 hours were discarded to remove artifacts from the day 1 recording. Amplitude and phase were identified by screening for the data point with maximum or minimum bioluminescence counts every 24 hours in RStudio and were checked by humans. The time of the first peak was determined as circadian phase, and the difference of bioluminescence counts in a cycle were calculated as the amplitude.

### EEG sleep recording

The same cohort of WT mice were surgically implanted with electrodes and habituated to TRL, LiFE and control schedules sequentially after recovery from surgery. We collected data for 24 hours on each experimental schedule. EEG and electromyography (EMG) signals derived from the surgically implanted electrodes were amplified and digitized at 256 Hz (Pinnacle). The signals were digitally filtered and spectrally analyzed by fast Fourier transformation, and polysomnographic recordings were scored using sleep analysis software (Sirenia Sleep Pro, Pinnacle). All scoring was performed manually based on the visual signature of the EEG and EMG waveforms, as well as the power spectra of 5 s epochs.

We defined wakefulness as desynchronized low-amplitude EEG and heightened tonic EMG activity with phasic bursts. We defined NREM sleep as synchronized, high-amplitude, low-frequency (0.5-4 Hz) EEG and highly reduced EMG activity compared with wakefulness with no phasic bursts. We defined REM sleep as having a pronounced theta rhythm (4-10 Hz) and a flat EMG. Microawakenings were defined as brief (<5 s) signatures of wakefulness in the EEG and EMG during NREM sleep. All sleep scoring was performed by investigators blind to the viral transgene delivered to the animal. Spectral analysis of EEG was performed by fast Fourier transform and binned in 0.5 Hz resolution from 0-25 Hz. Microawakening events were omitted from power analysis of NREM sleep.

### Food intake and bodyweight measurement

At the start of the experiment (light-off time), all food was removed from the cage top, and a few pellets (>20 grams) of standard diet were weighed and put to the metal cage top. For the food intake curve, every 4 hours during the next 24 hours, food remaining was weighed to calculate the food consumed. For daily food intake, 24 hours after the start of the experiment, all food remaining was weighed to calculate the food consumed. For daily food intake, measurements were repeated 3 days to calculate the average. Body weights of mice were measured by electronic scale at the time of light-off.

### Blood glucose measurement and glucose tolerance test (GTT)

Mice received a tail snip and blood glucose measure using a Glucometer (OneTouch Ultra Test Strips for Diabetes). For the baseline glucose curve, mice habituated to control, TRL, or LiFE treatment were measured at 0, 4, 8, 12, 16, and 20 hours after light-off.

In GTT, WT mice were overnight fasted for 20 hours prior to experiment start(+4 hours after light-off). Baseline fasted blood glucose was measured 15 minutes prior to dextrose (D-glucose) injection. At time point 0, mice received a blood glucose measure and injection of dextrose (1g/kg). At 15, 30, 60, and 90minutes after injection, blood glucose levels were measured. All measurements during the dark phase were taken in dark with a red light headlight to avoid circadian disruption.

### Morris water maze

Mice were habituated to the room of experiment for >30 mins. Experiments were conducted at 0-3 hours before light-off, with food provided and wheel locked for animals on LiFE schedule before start of experiment. Experiment procedures are modified based on published methods (58). The maze consisted of a circular pool (110 cm diameter, 25 cm height) filled with opaque water (∼21°C) made by adding non-toxic white paint. A circular escape platform (10 cm diameter) was positioned in a fixed location within the target quadrant (SW) throughout training. Distal visual cues of varying shapes and colors were fixed on the north, east, west and south corner outside of the pool wall and remained constant across all sessions. On day 1, mice were released at 4 starting points (N, E, NW, SE) with platforms 1-2 cm above the water. Animals were allowed for up to 60 seconds of exploration and guided to rest on the platform for 15 seconds. On days 2-5, the platform was hidden 1-2 cm below water and mice were released from the 4 starting points in a designed order that varied across days and were allowed for up to 60 s of exploration before guided to rest on the platform for 15 s. On day 6, the platform was removed from the pool and all animals were released from the corner furthest from the platform (NE) for free exploration of 60 s. Video of behaviors were acquired with the Ethovision system to analyze swim speed and time spent in each quadrant. Latency to the platform was manually recorded by stop watch.

### Novel object recognition

Mice were habituated in the room of experiment for >30 mins and experiments were conducted 0-3 hours before light-off with food provided and wheel locked for animals on LiFE schedule before start of experiment. A square arena (40 cm * 40 cm square * 10 cm height) with white floor and opaque walls was exposed to ∼ 150 lux illumination. Before each trial, the arena was sanitized with 70% ethanol to remove all scents. On the habituation day, animals spent 10 mins for free exploration in the empty arena. On day 1, mice freely explore in the arena with two identical objects positioned on the left and right side of the arena for 10 mins. 24 hours after the experiment on day 1, mice were placed back to the arena with the familiar object on the left side replaced with a novel object for 10 mins of free exploration. Videos were acquired and analyzed with the Ethovision system and software. Interactions with objects were determined as nose tracking of the animals within 2cm distance to the object. Preference of the object (right side) was calculated for both days as interaction time(right)/total interaction time. We defined the recognition index as preference (day 2) - preference (day 1).

### Alzheimer’s disease histology

Mice were sacrificed with and perfused immediately with PBS followed by 4% paraformaldehyde (PFA). Brains were extracted and fixed in 4% PFA for 24 hours, incubated in 30% sucrose in PBS for 2 days and stored in -80°C. Frozen brains were cryo-sectioned to 30 micron-thickness sections and stored in PBS. In immunohistochemistry, brain sections were rinsed in TBS buffer and incubated in blocking solution (Li COR cat# NC1660553 + 0.1% Tween-20) for 1 hour at room temperature (RT). Primary antibodies (D54D2, 1:2000, Cell Signaling cat# 8243T; AT8, Invitrogen 1:1000, cat# MN1020) were diluted in blocking solution and incubated with brain sections at 4℃ overnight. Brain sections were washed by TBST(TBS + 0.1% Tween-20) for 10 min for 3 times, and incubated with secondary antibodies (Invitrogen, Alexa 488 goat anti-rabbit 1:500, Alexa 647 goat anti-mouse 1:500) for 2 hours at room temperature. After secondary antibody incubation, brain sections were washed and mounted to glass slides. DAPI mounting medium (Southern Biotech cat# 0100-20) was used to cover the brain sections and the slide was sealed with cover slips for imaging.

### Confocal microscopy and histology image analysis

Mounted sections were imaged with 10X objective lens by Leica Stellaris confocal microscope. Images were captured with a frame sequence of each channel as an individual frame. Images are of 8-bit 4096*4096 resolution with a z stack of 10 μm step to generate a maximal projection that demonstrates histology staining covering the complete z axis of the imaged brain section.

Images were uploaded to ImageJ (FIJI) and subtracted background (rolling-ball radius = 50). The areas of interest were manually circled according to the Allen Institute mouse brain atlas. Images were transformed into a binary image with a fixed thresholding of 75/255. Area fractions of staining were measured with Fiji software and exported for analysis. For each mouse, 3 images were obtained from different brain sections as replicates.

### Statistics

All statistical comparisons were made using the appropriate test in RStudio or GraphPad Prism 9.5.0. Data are presented as individual data points and/or mean ± SEM. Data were tested for outliers using the Grubbs’ test, and statistical outliers were included in data analysis since the inclusion did not change statistical conclusion. Grouped analyses were tested by t-test, Browns-Forsythe test (data set with unequal standard deviations), Kolmogorov–Smirnov test (for distribution) or one-way ANOVA with Bonferroni test for multiple comparisons in GraphPad Prism or RStudio. Data measuring time, animal subject and measurement components were tested by linear mixed-effect model (time, experiment group and interaction as fixed effects and mouse ID as a random intercept) in R. Significance of fixed effects was evaluated using Type III ANOVA with Bonferroni post-hoc test.

## Data availability

All data are available on request.

## Acknowledgements

We thank all current and past members of the Güler lab and Bloom lab, especially Qijun Tang, Elizabeth N. Godschall, Taha Bugra Gungul, Isabelle R. Sajonia, O. Yipkin Calhan, Weili Liu, Jiachen Shi, Xuehan Sun, Andres Norambuena, Stefani Mancuso and Dora Bigler Wang. The work was supported by NIH R35GM140854 (ADG), Alzheimer’s Association AARG-NTF-21-852299 (ADG), University of Virginia Brain Institute Presidential Fellowship in Collaborative Neuroscience (YS).

## Author contributions

YS, GSB, ADG designed the research. YS, SDR, JTS, WAM, SNM, RS, EGM, CMP, LRG, SSK performed the experiments and processed data. YS, MEC, GSB, ADG did formal analysis on the data. YS, MEC, GSB, ADG wrote the paper.

---

**Figure S1.**
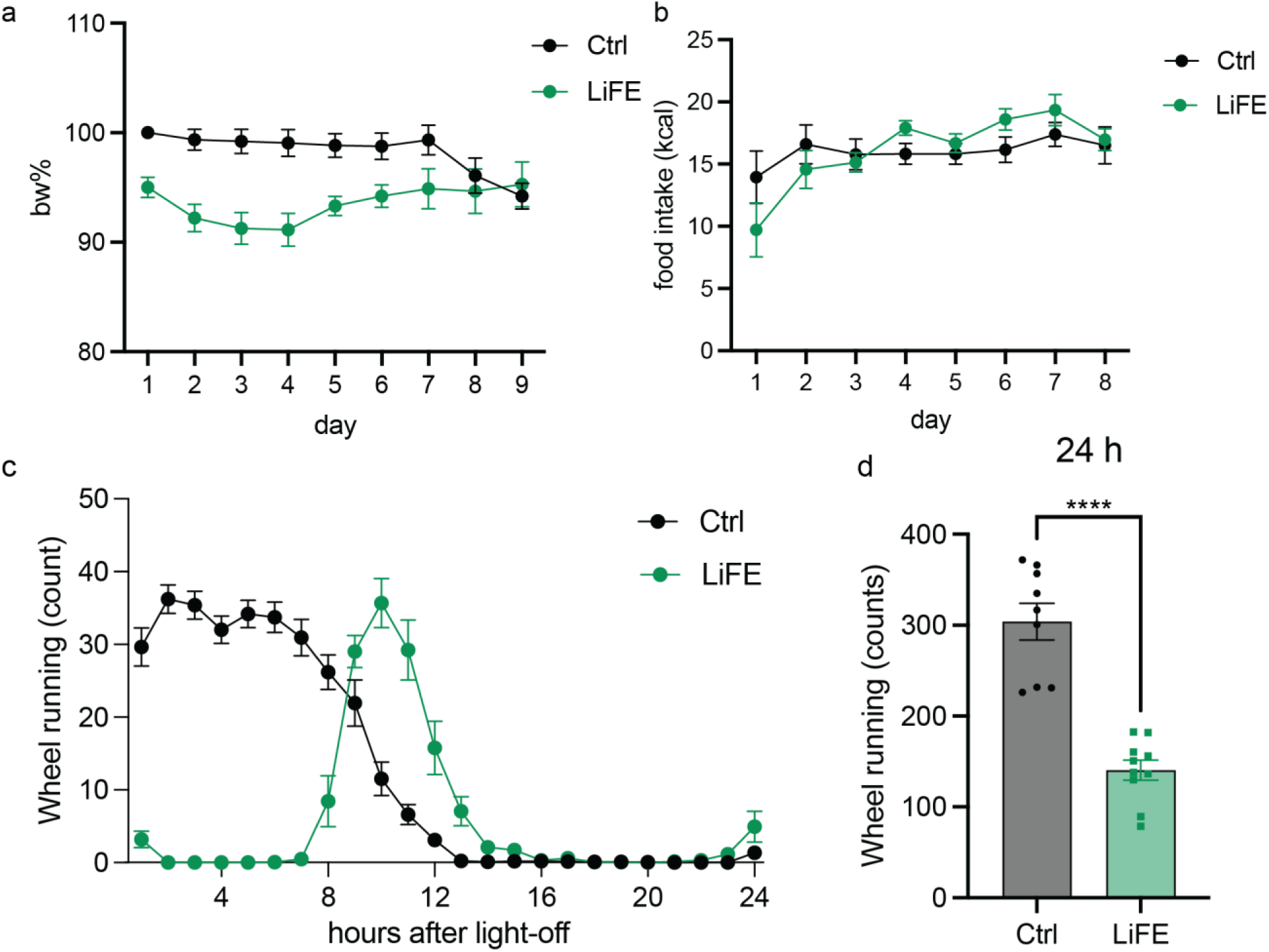
Characterizing behavior of WT mice in LiFE treatment. **a)** Bodyweight of WT mice on control (n = 10) and LiFE (n = 16) schedules normalized to the control weight of day 1. **b)** Food consumption in every 24 hours measured with WT mice (n =6) on control and LiFE schedule. **c)** Wheel running activity in a 1 hour bin of WT mice under control (n = 9) and LiFE (n = 10) schedule. **d)** Total wheel running counts of data in a) (Student’s t-test, \*\*\*\**P*<0.0001).

**Figure S2.**
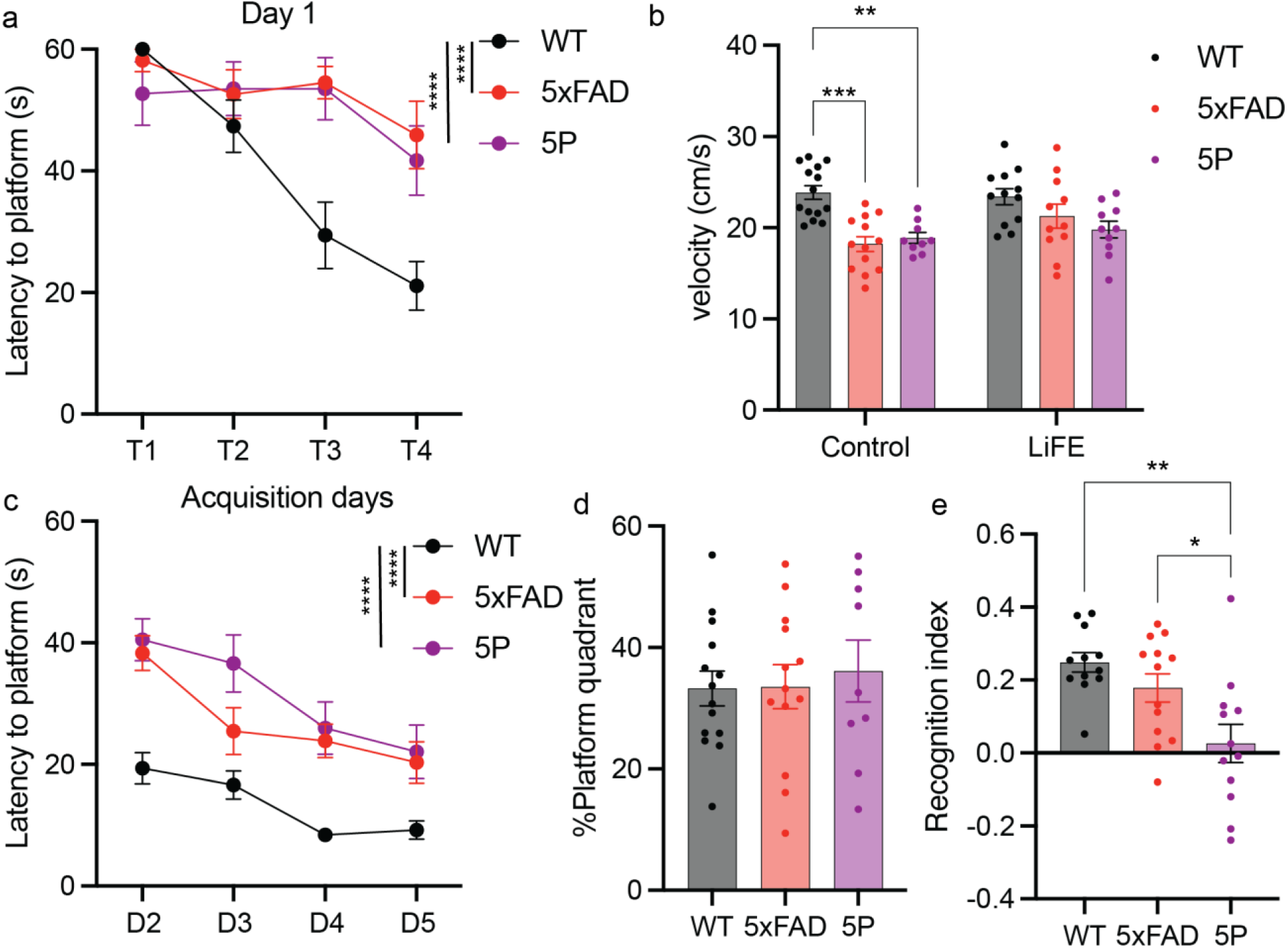
Comparison of cognitive behavior in WT and Alzheimer’s mouse models. **a)** Latency to the platform of WT (n = 14), 5xFAD (n = 13) and 5P (n = 9) control mice on day 1 learning in MWM (mixed-effect model with Bonferroni correction, \*\*\*\**P*<0.0001). **b)** Swim speed of WT (control n = 14, LiFE n = 12), 5xFAD (control n = 13, LiFE n = 11) and 5P (control n = 9, LiFE n = 10) mice in MWM test in control and LiFE treatment group (two-way ANOVA with Bonferroni correction, \*\**P*<0.01, \*\*\**P*<0.001).**c)** Latency to the platform of WT, 5xFAD and 5P control mice on acquisition days in MWM (mixed-effect model with Bonferroni correction, \*\*\*\**P*<0.0001). **d)** Time-spent in target quadrant of WT, 5xFAD and 5P control mice on probe day in MWM. **e)** Recognition index of WT (n = 12), 5xFAD (n = 13) and 5P (n = 12) control mice in NOR (one-way ANOVA with Bonferroni correction, \**P*<0.05, \*\**P*<0.01).

## References

1. J. S. Takahashi, Transcriptional architecture of the mammalian circadian clock. Nat. Rev. Genet. 18, 164–179 (2017).

2. S. M. Reppert, D. R. Weaver, Coordination of circadian timing in mammals. Nature 418, 935–941 (2002).

3. D. M. Berson, F. A. Dunn, M. Takao, Phototransduction by retinal ganglion cells that set the circadian clock. Science 295, 1070–1073 (2002).

4. D. M. Edgar, W. C. Dement, Regularly scheduled voluntary exercise synchronizes the mouse circadian clock. Am. J. Physiol. 261, R928–33 (1991).

5. M. E. Yurgel, et al., A stress-sensing circuit signals to the central pacemaker to reprogram circadian rhythms. Sci. Adv. 11, eadr7960 (2025).

6. R. Refinetti, Entrainment of circadian rhythm by ambient temperature cycles in mice. J. Biol. Rhythms 25, 247–256 (2010).

7. A. J. Davidson, F. K. Stephan, Feeding-entrained circadian rhythms in hypophysectomized rats with suprachiasmatic nucleus lesions. Am. J. Physiol. 277, R1376–84 (1999).

8. K. A. Stokkan, S. Yamazaki, H. Tei, Y. Sakaki, M. Menaker, Entrainment of the circadian clock in the liver by feeding. Science 291, 490–493 (2001).

9. B. Lee, J. Shao, Adiponectin and lipid metabolism in skeletal muscle. Acta Pharmaceutica Sinica B 2, 335–340 (2012).

10. Q. Tang, et al., Leptin receptor neurons in the dorsomedial hypothalamus input to the circadian feeding network. Sci. Adv. 9, eadh9570 (2023).

11. A. T. L. Hughes, et al., Timed daily exercise remodels circadian rhythms in mice. Commun. Biol. 4, 761 (2021).

12. S. Deota, et al., Diurnal transcriptome landscape of a multi-tissue response to time-restricted feeding in mammals. Cell Metab. 35, 150–165.e4 (2023).

13. A. Chaix, T. Lin, H. D. Le, M. W. Chang, S. Panda, Time-Restricted Feeding Prevents Obesity and Metabolic Syndrome in Mice Lacking a Circadian Clock. Cell Metab. 29, 303–319.e4 (2019).

14. E. F. Sutton, et al., Early Time-Restricted Feeding Improves Insulin Sensitivity, Blood Pressure, and Oxidative Stress Even without Weight Loss in Men with Prediabetes. Cell Metab. 27, 1212–1221.e3 (2018).

15. A. M. Schroeder, et al., Voluntary scheduled exercise alters diurnal rhythms of behaviour, physiology and gene expression in wild-type and vasoactive intestinal peptide-deficient mice. J Physiol (Lond) 590, 6213–6226 (2012).

16. A. Ramkisoensing, J. H. Meijer, Synchronization of biological clock neurons by light and peripheral feedback systems promotes circadian rhythms and health. Front. Neurol. 6, 128 (2015).

17. A. Nandi, et al., Cost of care for Alzheimer’s disease and related dementias in the United States: 2016 to 2060. npj Aging 10, 13 (2024).

18. E. S. Musiek, D. D. Xiong, D. M. Holtzman, Sleep, circadian rhythms, and the pathogenesis of Alzheimer disease. Exp. Mol. Med. 47, e148 (2015).

19. Y.-E. S. Ju, B. P. Lucey, D. M. Holtzman, Sleep and Alzheimer disease pathology--a bidirectional relationship. Nat. Rev. Neurol. 10, 115–119 (2014).

20. H. C. Hollis, et al., Reconstructed cell-type-specific rhythms in human brain link Alzheimer’s pathology, circadian stress, and ribosomal disruption. Neuron 113, 2822–2838.e7 (2025).

21. K. Madamanchi, J. Zhang, G. C. Melkani, Linkage of circadian rhythm disruptions with Alzheimer’s disease and therapeutic interventions. Acta Pharm. Sin. B 15, 2945–2965 (2025).

22. R.-Y. Pan, et al., Intermittent fasting protects against Alzheimer’s disease in mice by altering metabolism through remodeling of the gut microbiota. Nat. Aging 2, 1024–1039 (2022).

23. D. S. Whittaker, et al., Circadian modulation by time-restricted feeding rescues brain pathology and improves memory in mouse models of Alzheimer’s disease. Cell Metab. 35, 1704–1721.e6 (2023).

24. S. Fulda, et al., Rapid eye movements during sleep in mice: high trait-like stability qualifies rapid eye movement density for characterization of phenotypic variation in sleep patterns of rodents. BMC Neurosci. 12, 110 (2011).

25. M. Ezpeleta, et al., Time-restricted eating: Watching the clock to treat obesity. Cell Metab. 36, 301–314 (2024).

26. R. Stickgold, Sleep-dependent memory consolidation. Nature 437, 1272–1278 (2005).

27. T. A. LeGates, et al., Aberrant light directly impairs mood and learning through melanopsin-expressing neurons. Nature 491, 594–598 (2012).

28. E. S. Musiek, et al., Circadian Rest-Activity Pattern Changes in Aging and Preclinical Alzheimer Disease. JAMA Neurol. 75, 582–590 (2018).

29. S.-S. Yoo, P. T. Hu, N. Gujar, F. A. Jolesz, M. P. Walker, A deficit in the ability to form new human memories without sleep. Nat. Neurosci. 10, 385–392 (2007).

30. A. Saul, F. Sprenger, T. A. Bayer, O. Wirths, Accelerated tau pathology with synaptic and neuronal loss in a novel triple transgenic mouse model of Alzheimer’s disease. Neurobiol. Aging 34, 2564–2573 (2013).

31. H. Oakley, et al., Intraneuronal beta-amyloid aggregates, neurodegeneration, and neuron loss in transgenic mice with five familial Alzheimer’s disease mutations: potential factors in amyloid plaque formation. J. Neurosci. 26, 10129–10140 (2006).

32. S. D. Girard, et al., Onset of hippocampus-dependent memory impairments in 5XFAD transgenic mouse model of Alzheimer’s disease. Hippocampus 24, 762–772 (2014).

33. S. Jawhar, A. Trawicka, C. Jenneckens, T. A. Bayer, O. Wirths, Motor deficits, neuron loss, and reduced anxiety coinciding with axonal degeneration and intraneuronal Aβ aggregation in the 5XFAD mouse model of Alzheimer’s disease. Neurobiol. Aging 33, 196.e29–40 (2012).

34. S. Forner, et al., Systematic Phenotyping and Characterization of the 5xFAD mouse model of Alzheimer’s Disease. BioRxiv (2021) 10.1101/2021.02.17.431716.

35. . Y. Yoshiyama, et al., Synapse loss and microglial activation precede tangles in a P301S tauopathy mouse model. Neuron 53, 337–351 (2007).

36. . Y. Lee, J. M. Field, A. Sehgal, Circadian rhythms, disease and chronotherapy. J. Biol. Rhythms 36, 503–531 (2021).

37. . K. S. Bell, et al., Time-restricted feeding in adult mice improves mood-related behaviors in a sex-dependent manner. Behav. Brain Res. 496, 115867 (2026).

38. . V. D. Longo, S. Panda, Fasting, Circadian Rhythms, and Time-Restricted Feeding in Healthy Lifespan. Cell Metab. 23, 1048–1059 (2016).

39. . R. F. L. Vieira, et al., Time-restricted feeding combined with aerobic exercise training can prevent weight gain and improve metabolic disorders in mice fed a high-fat diet. J Physiol (Lond) 600, 797–813 (2022).

40. . Z. Dai, et al., The Effect of Time-Restricted Eating Combined with Exercise on Body Composition and Metabolic Health: A Systematic Review and Meta-Analysis. Adv. Nutr. 15, 100262 (2024).

41. . J. Liu, et al., Muscle Rev-erb controls time-dependent adaptations to chronic exercise in mice. Nat. Commun. 16, 5708 (2025).

42. . S. Tsameret, N. Chapnik, O. Froy, Effect of early vs. late time-restricted high-fat feeding on circadian metabolism and weight loss in obese mice. Cell. Mol. Life Sci. 80, 180 (2023).

43. . M. Sethi, et al., Increased fragmentation of sleep-wake cycles in the 5XFAD mouse model of Alzheimer’s disease. Neuroscience 290, 80–89 (2015).

44. . J. K. Holth, T. E. Mahan, G. O. Robinson, A. Rocha, D. M. Holtzman, Altered sleep and EEG power in the P301S Tau transgenic mouse model. Ann. Clin. Transl. Neurol. 4, 180–190 (2017).

45. . M. E. Wimmer, et al., Aging in mice reduces the ability to sustain sleep/wake states. PLoS ONE 8, e81880 (2013).

46. . J.-E. Kang, et al., Amyloid-beta dynamics are regulated by orexin and the sleep-wake cycle. Science 326, 1005–1007 (2009).

47. . M. Hatori, et al., Time-restricted feeding without reducing caloric intake prevents metabolic diseases in mice fed a high-fat diet. Cell Metab. 15, 848–860 (2012).

48. . L. M. Ince, et al., Time-restricted feeding rescues sociability deficits and reduces neuroinflammation in aged mice. Neurobiol. Aging 159, 1–14 (2026).

49. . K. Spiegel, E. Tasali, R. Leproult, E. Van Cauter, Effects of poor and short sleep on glucose metabolism and obesity risk. Nat. Rev. Endocrinol. 5, 253–261 (2009).

50. . H. T. VanderLeest, et al., Seasonal encoding by the circadian pacemaker of the SCN. Curr. Biol. 17, 468–473 (2007).

51. . T. Yoshikawa, et al., Localization of photoperiod responsive circadian oscillators in the mouse suprachiasmatic nucleus. Sci. Rep. 7, 8210 (2017).

52. . Y. Hu, et al., Voluntary wheel running exercise improves sleep disorder, circadian rhythm disturbance, and neuropathology in an animal model of Alzheimer’s disease. Alzheimers Dement 21, e70314 (2025).

53. . P. A. Adlard, V. M. Perreau, V. Pop, C. W. Cotman, Voluntary exercise decreases amyloid load in a transgenic model of Alzheimer’s disease. J. Neurosci. 25, 4217–4221 (2005).

54. . H. Song, et al., Aβ-induced degradation of BMAL1 and CBP leads to circadian rhythm disruption in Alzheimer’s disease. Mol. Neurodegener. 10, 13 (2015).

55. . J. Lee, et al., Inhibition of REV-ERBs stimulates microglial amyloid-beta clearance and reduces amyloid plaque deposition in the 5XFAD mouse model of Alzheimer’s disease. Aging Cell 19, e13078 (2020).

56. . M. W. King, S. M. Jacob, A. Sharma, J. H. Lawrence, E. S. Musiek, Circadian rhythms and the light-dark cycle interact to regulate amyloid-beta plaque accumulation and tau phosphorylation in 5xFAD mice. Alzheimers Dement 21, e70885 (2025).

57. . G. J. Kress, et al., Regulation of amyloid-β dynamics and pathology by the circadian clock. J. Exp. Med. 215, 1059–1068 (2018).

58. . C. V. Vorhees, M. T. Williams, Morris water maze: procedures for assessing spatial and related forms of learning and memory. Nat. Protoc. 1, 848–858 (2006).

